# uap: Reproducible and Robust HTS Data Analysis

**DOI:** 10.1101/690438

**Authors:** Christoph Kämpf, Michael Specht, Alexander Scholz, Sven-Holger Puppel, Gero Doose, Kristin Reiche, Jana Schor, Jörg Hackermüller

## Abstract

**Background:** A lack of reproducibility has been repeatedly criticized in computational research. High throughput sequencing (HTS) data analysis is a complex multi-step process. For most of the steps a range of bioinformatic tools is available and for most tools manifold parameters need to be set. Due to this complexity, HTS data analysis is particularly prone to reproducibility and consistency issues. We have defined four criteria that in our opinion ensure a minimal degree of reproducible research for HTS data analysis. A series of workflow management systems is available for assisting complex multi-step data analyses. However, to the best of our knowledge, none of the currently available work flow management systems satisfies all four criteria for reproducible HTS analysis.

**Results:** Here we present `uap`, a workflow management system dedicated to robust, consistent, and reproducible HTS data analysis. `uap` is optimized for the application to omics data, but can be easily extended to other complex analyses. It is available under the GNU GPL v3 license at https://github.com/yigbt/uap.

**Conclusions:** `uap` is a freely available tool that enables researchers to easily adhere to reproducible research principles for HTS data analyses.

## Background

Next generation or high throughput sequencing (HTS) methods that rely on massively parallel DNA sequencing have opened a new era of molecular life sciences. A continuous growth in sequencing throughput, precision, length of the sequencing reads, and increasing automation and miniaturization led to a huge advantage in manual efforts and costs per experiment compared to traditional Sanger sequencing. HTS has thus become a mainstay in many fields of biology and biomedicine: Apart from *de novo* genome sequencing, HTS is not only increasingly replacing other massively parallel techniques like microarrays in transcriptomics or epigenomics, but also allows for new approaches e.g. in microbial ecology, population genetics, clinical diagnostics, or breeding. As a consequence, the amounts of HTS-generated data are exploding in many disciplines, associated with challenges for storage, transfer, curation and particularly reproducible analysis of these data [1].

HTS data analysis is a multi-step process and fairly complex compared to the analysis of other biological data. Analysis of e.g. gene expression microarray data is involving image processing and statistical analysis of the expression signal. Analysis of a comparable transcriptome sequencing (RNA-seq) data set involves – apart from the initial base calling – at least clipping of adapter and low quality sequences, mapping to a reference genome, counting of mapped reads per annotation element and statistical analysis and may include many other steps like transcript *de novo* assembly or analysis of differential splicing. For most of these steps a selection from a range of available bioinformatic tools needs to be made and for most tools a variety of parameters needs to be set.

In general, an HTS data analysis can be described as a directed acyclic graph (DAG) like structure. We call a node in the DAG a step. Steps may depend on previously computed results (produced by preceding steps) and/or branch out to subsequent parts of the analysis. Results of individual steps also may be merged again in later steps and processed further. See Figure 1 for a sketch of a prototypic HTS analysis.

**Figure 1.**
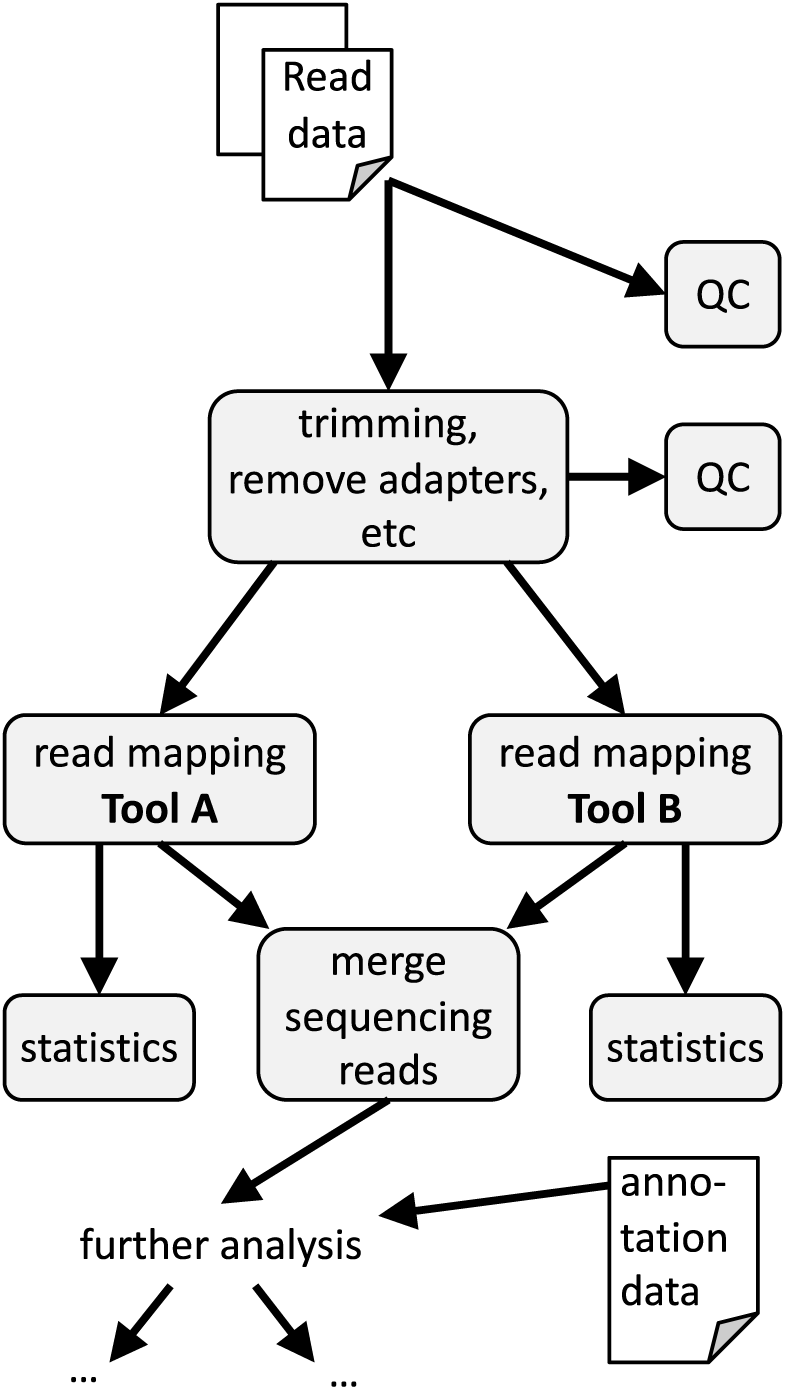
Sketch of a prototypic DAG describing the analysis of unmapped HTS reads. HTS data analysis typically follows a DAG-like structure. Nodes of the DAG are called steps and may depend on preceding steps and/or branch out to subsequent parts of the analysis. Results of individual steps may be merged again in later steps and processed further.

Independent replication is a fundamental principle for evaluating published findings. If a complete replication is not feasible e.g. due to the access to samples or cost and effort for HTS experiments, reproducing the analysis from raw data to published claims is the second best alternative [2]. The degree of reproducibility of biomedical research has been criticized [3] and a “reproducibility crisis” has been diagnosed recently [4]. Given the complexity of HTS data analyses outlined above, reproducibility is particularly dependent on a detailed reporting of analysis details. However, a critical analysis of published HTS-based genotyping studies revealed that less than a third of the studies analyzed provided sufficient information to reproduce the mapping step [1].

Several appeals have been made to alleviate these reproducibility issues in computer science and computational biology. Roger Peng emphasized the necessity of linking executable analysis code and data as the gold standard second to full replication [2]. Sandve and colleagues called for adhering to “ten simple rules for reproducible computational research”, which fully apply to HTS data analysis [5]. Finally, Grüning and coauthors defined a technology stack for reproducible research and formulate guidelines that particularly consider the numerical reproducibility of computation in the life sciences [6].

In our opinion, a minimal degree of reproducible research in managing HTS data analyses requires a tool which ensures that (*i*) the dependencies between analysis steps and intermediate results are correctly maintained, (*ii*) analysis steps are successfully completed prior to execution of subsequent steps, (*iii*) all tools, their versions and full parameter sets (including standard parameters which are usually not set when starting the tool from the commandline) are logged, (*iv*), the consistency between the code defining the analysis and the currently available results is ensured.

A series of different bioinformatic workflow management systems (WMS) is available to support complex DAG-like analyses. WMS that are appropriate for HTS analyses are in part general purpose systems, in part specific for HTS or even designed to address individual aspects of HTS data analysis. Also, different WMS designs require various levels of experience from the users, while providing different degrees of flexibility. WMS approaches like `Ruffus` [7] or `SnakeMake` [8] allow the implementation of highly individual analyses, either via domain-specific programming languages or a general-purpose programming language. `iRAP` [9], `RseqFlow` [10], or `MAP-RSeq` [11] belong to a group of WMS that implement a single specific type of HTS analysis. Several WMS encapsulate the individual steps of an analysis within modules and allow for their free combination, e.g. `Galaxy` [12], `Unipro UGENE` [13], `KNIME` [14], or `Taverna` [15] – many of which come with a graphical user interface. Finally, a group of lightweight modularized WMS aims at a modular, customizable, command-line-based approach, which includes e.g. `bcbio-nextgen` [16], `Bpipe` [17], or `nextflow` [18]. Several of these WMS rely on logging detailed information of used tools’ versions and issued commands. However, to the best of our knowledge, none of the published WMS satisfies all four criteria that we defined for maintaining a minimal degree of reproducible research in HTS data analysis. We compared the essential features, particularly regarding reproducibility, of a set of actively maintained, flexible, and modular WMS in more detail (Table 1, Supplemental Table S 1.

**Table 1.**
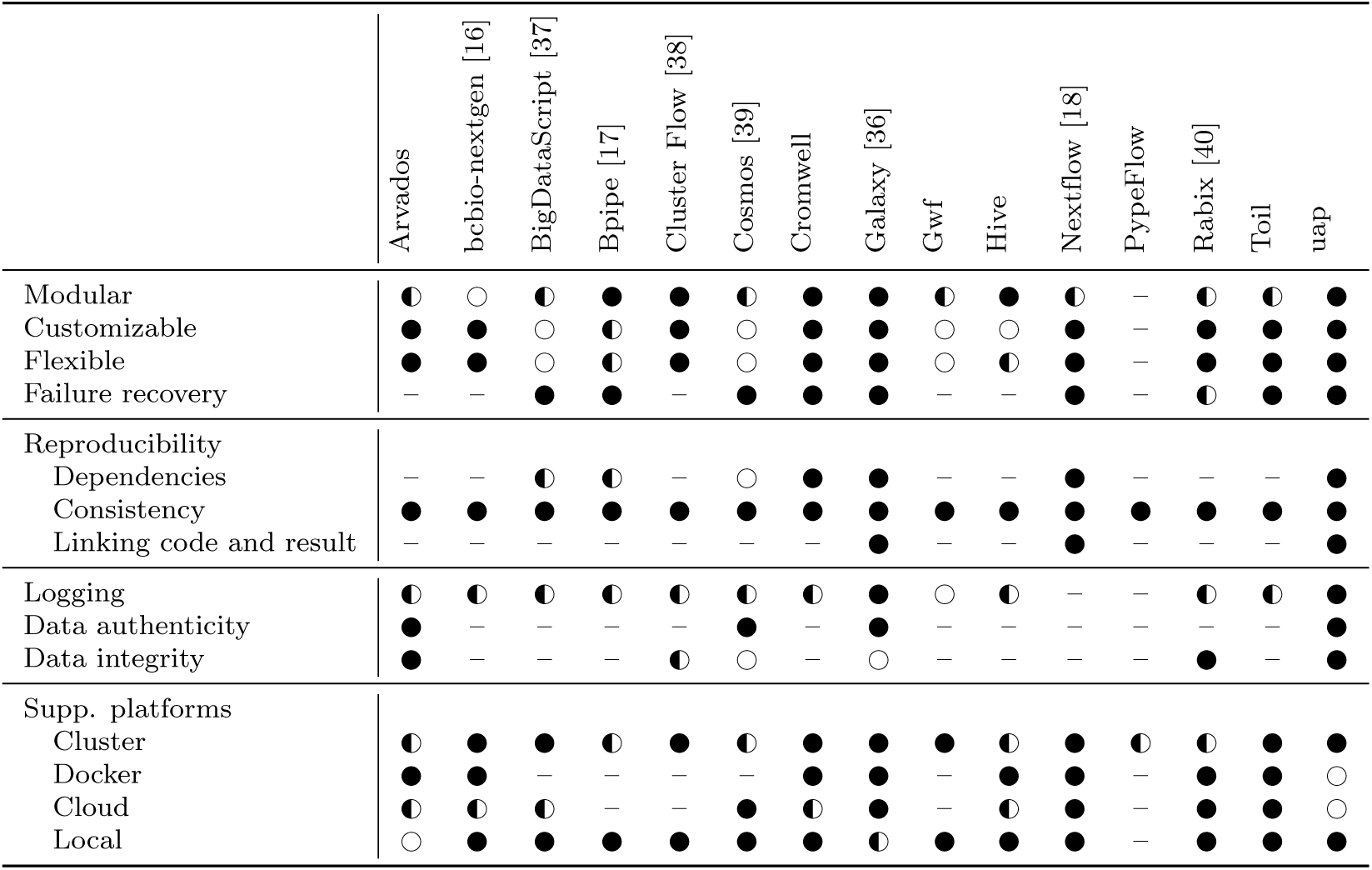
Comparison of workflow management systems and HTS analysis pipelines based on essential features, particularly regarding reproducibility. Tools were selected from https://github.com/pditommaso/awesome-pipeline retaining only tools that were considered to be actively developed by at least five contributors (latest commit after 31.01.2018), are licensed as open source, where the scope of application is clearly on bioinformatics, and which support cluster batch systems. **Modular**: Workflows are assembled from reusable steps; **Customizable**: Configuration of the analysis is separated from the code defining the steps; **Flexible**: Workflows can easily be altered without programming skills; **Failure recovery**: Failure during execution is detected and subsequent analyses halted to avoid corrupted results; **Reproducibility - Dependencies**: WMS enforces that dependencies between analysis steps and intermediate results are correctly maintained. **Consistency**: WMS safeguards that analysis steps are successfully completed prior to execution of subsequent steps. **Linking code and result**: WMS ensures consistency between the code defining the analysis and the currently available results. **Logging**: WMS logs Stdout/Stderr, exit status, tool versions, WMS version, executed commands, execution date, and in-/output files (see corresponding supplemental table for details); **Data authenticity** WMS records information about creator and creating process(es) of the data; **Data integrity**: WMS records information that allows to verify the integrity of created data (e.g. hash sums); **Supp. platforms - Cluster**: WMS supports compute cluster management systems ton run jobs; **Docker**: WMS can run jobs using Docker images; **Cloud**: WMS can be deployed in a compute cloud; **Supp. CWL**: WMS supports common workflow language; **Local**: WMS can be executed locally without depending on a cluster management system or a (web)server; Results are given as ○ (not met), ◐ (partially met), ● (fully met) and – (not stated). Ratings are based on information provided in the papers, documentations and manuals. A more detailed comparison is presented in Supplemental Table S 1.

Here, we introduce the workflow management system `uap` (Universal Analysis Pipeline) that may be used to implement any DAG-like data analysis workflow, but is primarily aimed at HTS data analysis. It is constructed to execute, control, and keep track of the analysis of large data sets. uap encapsulates the usage of (not necessarily bioinformatic) tools and handles data flow and processing of the complete analysis. Produced data is tightly linked with the code specifying the analysis. Thus, it enables users to perform reproducible, robust, and consistent data analyses. We provide complete workflows for handling genomic data and analyzing RNA-Seq and Chromatin Immuno-Precipitation DNA-Sequencing (ChIP-seq) data, which can be used as templates that allow for easy customization. As we are also integrating steps for downloading published sequencing raw data (e.g. from SRA), `uap` enables users to efficiently reproduce the data analysis of published studies. The provided workflows have been optimized for minimal I/O load on high performance computing (HPC) environments. Although, initially designed for HTS data analysis, the plugin architecture of `uap` allows for the expansion to any kind of data analysis.

## Implementation

`uap` is a workflow management software (WMS) implemented in Python. It provides user-friendly access to a range of bioinformatic analyses of large datasets, such as high throughput sequencing data. Each analysis is completely described by an individual configuration file in YAML^1^ format. The steps of the analysis and their dependencies as well as the required tools, parametrization and locations in the file system are specified there. Based on these settings `uap` constructs a directed acyclic graph (DAG) that represents the workflow of the analysis.

### Analysis as a directed acyclic graph

The DAG represents single analysis steps as nodes and pairwise dependencies between steps as directed edges. A *step* is a blueprint for a particular analysis with defined input and output data. The passing of input data to a step and the generation of a particular type of output data is modeled as *connections*. They control the data flow between steps by grouping output files and providing them to down-stream steps. `uap` distinguishes between source and processing steps. *Source steps* emit data into the workflow and hand the files over to downstream processing steps via output connections. The user is free to categorize the input data files for a workflow into user specified groups to create separate output locations for each category. *Processing steps*, on the other hand, receive data from upstream steps via input connections, define a sequence of execution commands and assign output file locations for each input connection. The entirety of these configurations of a step for a particular set of input connections is called a *run*. It can be interpreted as an instance of a step and is the atomic unit of the analysis. Supplemental Figure S 1 shows the DAG including its runs rendered by `uap` based on the configuration file for the analysis of a published data set.

### Plug-in architecture

Steps encapsulate the usage of a tool in a single python class, which allows users to easily customize `uap` by adding steps. Every new step inherits from a super class, defines incoming and outgoing connections, the required tools, and has to implement the `runs()` method. A step can individually be optimized for efficient use of CPU and memory usage. For allowing a flexible accommodation to different high performance computing environments, `uap` supports a step-specific adaptation of the environment, e.g. for setting variables, or automatic loading and unloading of software modules.

### Enforcing consistency and integrity

When computing on large data sets, partial processing of large files due to premature termination of tools may remain undetected without stringent monitoring of processes and poses a severe threat to data integrity. In `uap`, runs are therefore executed in a temporary directory and monitored throughout execution. The overall workflow is not compromised in case a single run fails. Result files are only moved to their final location if all processes of a run exited gracefully and all expected output files exist.

`uap` automatically re-schedules runs if it detects failed processes or missing files. Also, changes in the configuration trigger re-scheduling of the affected runs and all dependent runs in the DAG.

### Maintaining reproducibility

`uap` tightly links analysis code and resulting data by hashing over the complete sequence of commands including parameter specifications of a run and appending the key to the output path. Thus, any changes to the analysis code alter the expected output location, which allows `uap` to check whether analysis code and output correspond to each other.

At execution time, an annotation file in YAML format is captured for each run that contains the complete content of the configuration at this point. Hence, an executed run is documented with the releases of all used software and the invoked command line with all parameter settings. In addition, memory and CPU usage of each process, checksums of result files, as well as the last kB of `stdout` and `stderr` output are reported. The annotation file is stored next to the result files of a run.

### Process flow

Initially, `uap` reads the configuration, generates the respective DAG, and defines all commands and output file names. Throughout this initiation process `uap` inspects the planned analysis for potential errors. The graph is tested to be acyclic, all required tools (in defined releases) are tested for their availability and the status of all steps is determined. This initiation phase is executed early, i.e. before submitting runs to a compute cluster. `uap` thus implements a failing fast technique. This is an important feature when working with large amounts of data on HPC systems where software is dynamically loaded and erroneous configurations might otherwise only become apparent after hours of computation. Figure 2 illustrates `uap`’s process flow, error reporting, and the link between configuration and result files.

**Figure 2.**
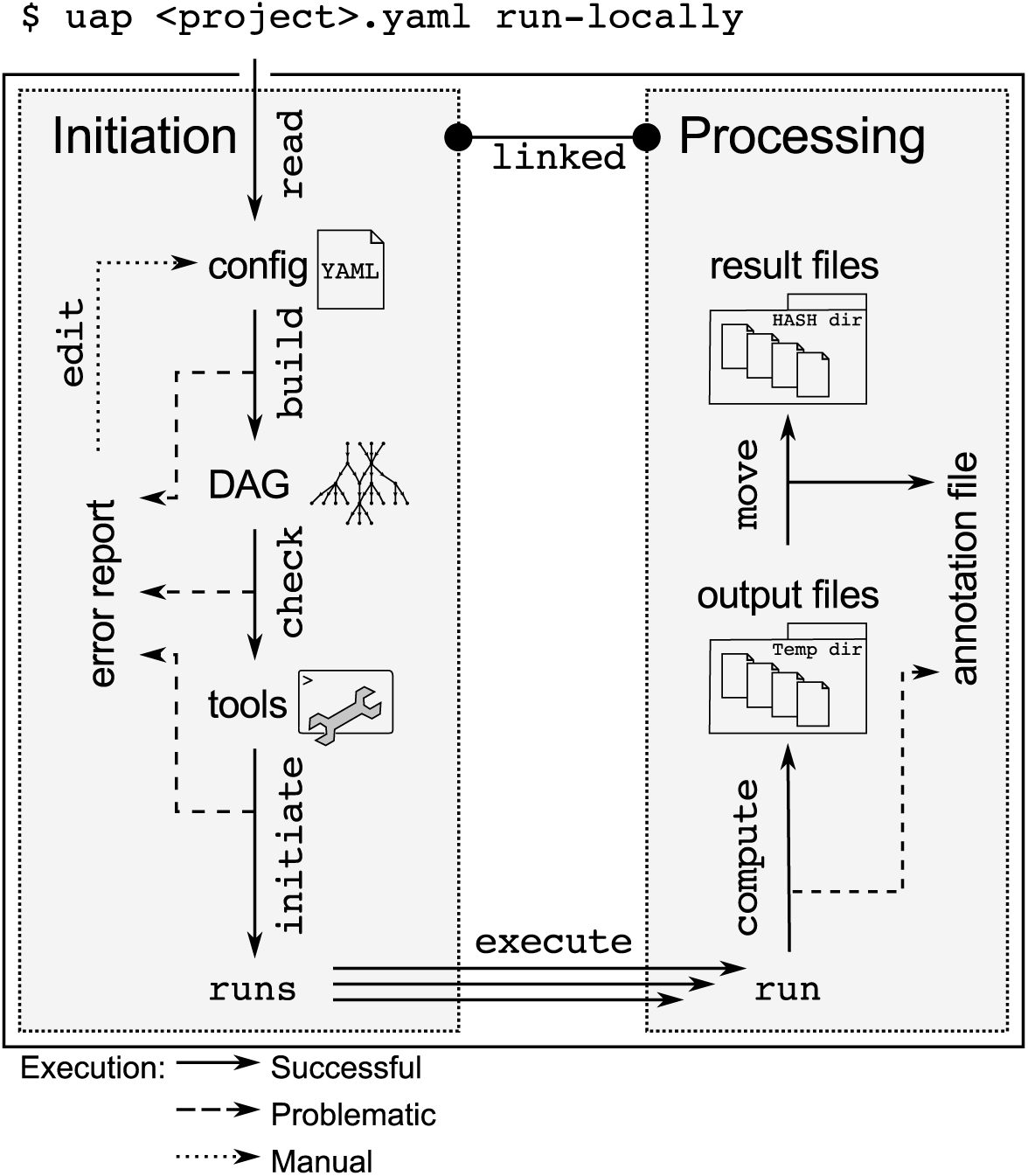
uap’s process flow, error reporting, and the link between the analysis code and result files. `uap` implements a failing fast approach: the DAG is built from the configuration file, tested to be acyclic, all required tools are tested for their availability and the status of all steps is determined. Subsequently, `uap` can start runs, display the commands of runs, show the state of the runs, and render execution graphs. Runs are executed in temporary directories and monitored throughout execution. Result files are only generated at their final location if all processes of a run exited gracefully and all expected output files exist. Analysis code and resulting data are tightly linked by hashing over the complete sequence of commands and parameters of a run and appending the key to the output path. Each run generates an annotation file in YAML format that captures the configuration, software versions and releases, the invoked command line, all parameters, memory and CPU usage of each process involved, checksums of the result files, as well as the last kB of `stdout` and `stderr`.

Subsequently, `uap` can start runs, display the commands of runs, show the state of the runs, and render execution graphs. Execution graphs are useful tools to inspect the performance of an analysis, e.g. to identify resource bottlenecks in a pipeline of commands. Supplemental Figure S 2 shows such an execution graph.

## Results

`uap` is a workflow management system dedicated to data consistency and adoption of a Reproducible Research paradigm in HTS data analysis. `uap` runs on UNIX-like operating systems and can interact with batch queuing systems like the Sun/Oracle/Univa grid engines (SGE/OGE/UGE) and SLURM [19] to submit analyses to high performance computing systems. `uap` is distributed under the GNU GPL v3 license and is publicly available at https://github.com/yigbt/uap. Its documentation is hosted at http://uap.readthedocs.org/. A `docker` container with a core set of tools is available at https://hub.docker.com/r/yigbt/uap/tags

`uap` is distributed with predefined workflows for (*i*) genome sequence download and index generation for read mapping programs, (*ii*) transcriptome sequencing (RNA-seq) data analysis, and (*iii*) Chromatin Immuno-Precipitation DNA-Sequencing (ChIP-Seq) data analysis. Further, we provide small test data sets enabling a quick start for each of these workflows. An additional example using a larger data set is provided via code for downloading and analyzing a publicly available ChIP-Seq data set (Barski *et al.* [20]). The provided workflows are intended to serve as an easy entry point into a `uap` analysis as well as a template for similar analyses, e.g. for other species or with another set of tools.

### Preparing genomic data for HTS analysis

An important prerequisite for HTS projects where a reference genome is available is aligning (mapping) the sequencing reads to this genome. Most mapping software requires a specific data structure (index) of the genome to efficiently solve this alignment problem. Indices have to be generated once, prior to any mapping procedure. We provide `uap` configuration files for the `bwa` [21], `bowtie` [22] and `segemehl` [23] mapping programs and `samtools` (`fasta` indexing) of the a) *Mycoplasma genitalium*, and b) *Homo sapiens* genome. Genomic sequences can be downloaded automatically prior to index generation.

### Transcriptome assembly from RNA-Seq data

Transcriptome sequencing identifies and estimates the quantity of RNA in biological samples. Beyond quantification of known transcripts based on overlapping sequence reads, RNA-seq allows the assembly of novel transcripts. We provide a `uap` configuration file for combining split-read mapping with *de novo* transcript assembly. uap reads the sequencing data either from an Illumina sequencing run folder, or a set of fastq files, applies quality control, removes adapter sequences, and maps the reads to a genome using `tophat2` [24] and `segemehl` [23, 25]. The mapped reads from `tophat2` are directly processed by `cufflinks` [26] for *de novo* transcript assembly. Split-reads mapped with `segemehl` are prepared for `cufflinks` using an adapter script and then also processed with `cufflinks`. The configuration also contains a step to determine the number of mapped reads per transcript applying `htseq-count` [27].

### Identification of enriched regions from ChIP-Seq data

ChIP sequencing is a method that integrates chromatin immunoprecipitation (ChIP) and high-throughput DNA sequencing to identify sites of protein-DNA interactions. The provided `uap`-configuration file for this task initially resembles the RNA-Seq workflow. Here, reads are expected to correspond to genomic DNA and the mapping is done without considering split-reads using `bowtie2` [22]. Mapped reads are subsequently sorted, duplicates removed, and enriched regions are detected using `MACS2` [28].

## Conclusions

The critique on a lack of reproducibility in science and a growing awareness that many reported facts do not seem to hold up to repeated investigation has meanwhile reached a broader audience beyond the scientific community (e.g. [29–31]). In our opinion, HTS data analysis is particularly prone to consistency and reproducibility issues – especially due to the complexity of the analysis, the involved data volume, and the broad range of available tools and their multitude of parameters.

Workflow management systems are indispensable for controlling more complex analyses, like HTS data analysis. Published workflow management systems for HTS data analysis are either highly flexible and made for experienced programmers or lack a lot of flexibility but can be used intuitively and some are specific to a certain type of analysis. In the introduction we listed four minimal requirements that we consider essential for ensuring reproducibility and consistency of HTS data analyses. None of the published systems we are aware of, however, completely satisfies these criteria. `uap` has been designed to fulfill these. One critical requirement is linking analysis code and generated data. While *Reproducible Research* in statistics [32] uses tools like Sweave [33], knitR [34], or Jupyter [35] to combine analysis code and resulting data in one output file, such a strategy is not feasible for most steps of an HTS analysis due to the size of generated data. `uap` therefore relies on hashing over the complete sequence of commands including parameters of a run and appending the key to the output path. In addition, `uap` performs logging and process monitoring, supports different cluster management systems, creates recovery points, plots execution graphs, manages job dependencies, and is extensible to any kind of multi-step analysis. It provides pre-built steps for the preparation of genomic data and the analysis of RNA-Seq and ChIP-Seq data.

Among the many different flavors of WMS `uap` is clearly easier to operate for users with limited programming experience than systems based on domain specific programming languages, while offering a lot more flexibility than single purpose tools. Based on the comparison of tools in Supplemental Table 1, the tools most dedicated to reproducible research and providing the most similar set of features compared to `uap` are `Galaxy` and `Nextflow`.

In our opinion, `Galaxy` [36] and `uap` address different user groups and tasks and we use both WMS in our research environment. `Galaxy` is great for providing predefined workflows to users without experience in programming and working on the command line. It allows these users to adapt parameters and execute such workflows on their data. For users working frequently with large HTS data sets, adapting workflows to a larger extent or duplicating branches of the DAG for performing variants of the analysis in parallel is much more efficiently performed in `uap`. `Galaxy` does not link data and code in the sense of `uap`. But as any change to parameters in a `Galaxy` workflow triggers re-execution of the sub-workflow below, this is not necessary. If many changes throughout a workflow have to be made, this behavior of `Galaxy` may be hindering. Obviously, running HTS analyses on `Galaxy` requires a `Galaxy` server integrated with an HPC environment, which is not trivial to set up and demands continuous maintenance. Starting from scratch, setting up HTS analysis is significantly less effort using `uap` than `Galaxy`.

`Nextflow` is a powerful WMS dedicated to scalability and reproducibility [18]. Its approach to reproducibility relies on a tight integration with `github` and the support of scalable containerization of pipelines using e.g. `Docker`. `Nextflow` and `uap` share several concepts, e.g. using temporary files for intermediate results or analyzing the workflow DAG to enable failing fast. Intermediate results of an HTS analysis can consume large amounts of storage space. `uap` therefore provides a means for volatilization of intermediate results without breaking dependencies in the DAG - a feature which does not seem to be available in `Nextflow`. Logging is somewhat limited in `Nextflow` compared to `uap`, but `Nextflow` provides a broader support for HPC environments including also support for cloud computing. `Nextflow’s` approach to reproducibility is powerful when software from `Github` is used, as it enables the user to request a specific commit, or when the tools used are publicly available as a container. However, when a tool is run in ‘native task support’ like the `Kallisto` example provided in [18], `uap` is more stringent in logging version and the full set (including default) of parameters. Where `uap` and `Nextflow` differ most clearly regarding reproducibility is linking data and code, as to our understanding based on publications and the online documentation this is not available in `Nextflow`.

In summary, we are convinced that reproducible research principles need to be advanced for HTS data analysis and that `uap` is a highly useful system for facing this challenge.

## Availability and requirements

**Project name:** uap

**Project home page:** https://github.com/yigbt/uap

**Operating system(s):** Linux.

**Programming language:** Python.

**Other requirements:** virtualenv, git, and graphviz.

**License:** GNU GPL v3.

**Any restrictions to use by non-academics:** None.

## Abbreviations

ChIP-seq: Chromatin Immuno-Precipitation DNA-Sequencing
DAG: directed acyclic graph
HTS: high throughput sequencing
HPC: high performance computing
RNA-seq: transcriptome sequencing
WMS: workflow management system

## Availability of data and material

Source code is available at https://github.com/yigbt/uap, links to Docker containers and documentation are provided at https://www.ufz.de/index.php?en=44919. All datasets used within the example workflows are publicly available and are fully referenced within the workflow configuration files.

## Author’s contributions

KR, MS, and JH initially designed and implemented the software. SHP, CK, AS, KR, JS, and JH took over the further development of the software. GD implemented the script that formats segemehl-output to be compatible with the cufflinks suite of tools. CK and AS provided the docker containers of uap. CK, KR, JS, and JH wrote the manuscript. All authors read, corrected and approved the final manuscript.

## Acknowledgments

This work was supported in part by the Initiative and Networking Fund of the Helmholtz Association through a grant to JH (VH-NG738), by the Deutsche Forschungsgemeinschaft (DFG) through a grant to JH (SPP1738), by the Fraunhofer Future Fund, and through the LIFE Leipzig Research Center for Civilization Diseases.

## Additional Files

**Additional file 1 — Supplemental_tables_and_figures.pdf**

This file includes supplemental tables and figures.

## Footnotes

1 http://yaml.org/

## References

1. Nekrutenko, A., Taylor, J.: Next-generation sequencing data interpretation: enhancing reproducibility and accessibility. Nature reviews. Genetics 13, 667–672 (2012). doi: 10.1038/nrg3305

2. Peng, R.D.: Reproducible research in computational science. Science (New York, N.Y.) 334, 1226–1227 (2011). doi: 10.1126/science.1213847

3. Bustin, S.A.: The reproducibility of biomedical research: Sleepers awake! Biomolecular detection and quantification 2, 35–42 (2014). doi: 10.1016/j.bdq.2015.01.002

4. Baker, M.: 1,500 scientists lift the lid on reproducibility. Nature 533, 452–454 (2016). doi: 10.1038/533452a

5. Sandve, G.K., Nekrutenko, A., Taylor, J., Hovig, E.: Ten simple rules for reproducible computational research. PLoS computational biology 9, 1003285 (2013). doi: 10.1371/journal.pcbi.1003285

6. Grüning, B., Chilton, J., Köster, J., Dale, R., Soranzo, N., van den Beek, M., Goecks, J., Backofen, R., Nekrutenko, A., Taylor, J.: Practical computational reproducibility in the life sciences. Cell systems 6, 631–635 (2018). doi: 10.1016/j.cels.2018.03.014

7. Goodstadt, L.: Ruffus: A lightweight python library for computational pipelines. Bioinformatics 26, 2778–2779 (2010). doi: 10.1093/bioinformatics/btq524

8. Köster, J., Rahmann, S.: Snakemake-a scalable bioinformatics workflow engine. Bioinformatics 28(19), 2520–2522 (2012). doi: 10.1093/bioinformatics/bts480

9. Fonseca, N.a., Petryszak, R., Marioni, J.: iRAP - an integrated RNA-seq Analysis Pipeline iRAP - an integrated RNA-seq Analysis Pipeline, 0-10 (2014). doi: 10.1101/005991

10. Wang, Y., Mehta, G., Mayani, R., Lu, J., Souaiaia, T., Chen, Y., Clark, A., Yoon, H.J., Wan, L., Evgrafov, O.V., Knowles, J.A., Deelman, E., Chen, T.: RseqFlow: Workflows for RNA-Seq data analysis. Bioinformatics 27(18), 2598–2600 (2011). doi: 10.1093/bioinformatics/btr441

11. Kalari, K.R., Nair, A.a., Bhavsar, J.D., O Brien, D.R., Davila, J.I., Bockol, M.a., Nie, J., Tang, X., Baheti, S., Doughty, J.B., Middha, S., Sicotte, H., Thompson, A.E., Asmann, Y.W., Kocher, J.-P.a.: MAP-RSeq: Mayo Analysis Pipeline for RNA sequencing. BMC bioinformatics 15(1), 224 (2014). doi: 10.1186/1471-2105-15-224

12. Goecks, J., Nekrutenko, A., Taylor, J.: Galaxy: a comprehensive approach for supporting accessible, reproducible, and transparent computational research in the life sciences. Genome biology 11, 86 (2010). doi: 10.1186/gb-2010-11-8-r86

13. Golosova, O., Henderson, R., Vaskin, Y., Gabrielian, A., Grekhov, G., Nagarajan, V., Oler, A.J., Quiñones, M., Hurt, D., Fursov, M., Huyen, Y.: Unipro UGENE NGS pipelines and components for variant calling, RNA-seq and ChIP-seq data analyses. PeerJ 2, 644 (2014). doi: 10.7717/peerj.644

14. Berthold, M.R., Cebron, N., Dill, F., Gabriel, T.R., Kötter, T., Meinl, T., Ohl, P., Sieb, C., Thiel, K., Wiswedel, B.: KNIME: The Konstanz Information Miner. In: Studies in Classification, Data Analysis, and Knowledge Organization (GfKL 2007). Springer, New York (2007)

15. Wolstencroft, K., Haines, R., Fellows, D., Williams, A., Withers, D., Owen, S., Soiland-Reyes, S., Dunlop, I., Nenadic, A., Fisher, P., Bhagat, J., Belhajjame, K., Bacall, F., Hardisty, A., Nieva de la Hidalga, A., Balcazar Vargas, M.P., Sufi, S., Goble, C.: The Taverna workflow suite: designing and executing workflows of Web Services on the desktop, web or in the cloud. Nucleic acids research 41(Web Server issue), 557–561 (2013). doi: 10.1093/nar/gkt328

16. Guimera, R.V.: bcbio-nextgen: Automated, distributed next-gen sequencing pipeline. EMBnet.journal 17, 30 (2012). doi: 10.14806/ej.17.B.286

17. Sadedin, S.P., Pope, B., Oshlack, A.: Bpipe: a tool for running and managing bioinformatics pipelines. Bioinformatics (Oxford, England) 28(11), 1525–6 (2012). doi: 10.1093/bioinformatics/bts167

18. Di Tommaso, P., Chatzou, M., Floden, E.W., Barja, P.P., Palumbo, E., Notredame, C.: Nextflow enables reproducible computational workflows. Nature biotechnology 35, 316–319 (2017). doi: 10.1038/nbt.3820

19. Yoo, A.B., Jette, M.A., Grondona, M.: SLURM: Simple Linux Utility for Resource Management. In: Feitelson, D., Rudolph, L., Schwiegelshohn, U. (eds.) Job Scheduling Strategies for Parallel Processing: 9th International Workshop, JSSPP 2003, Seattle, WA, USA, June 24, 2003. Revised Paper, pp. 44–60. Springer, Berlin, Heidelberg (2003). http://dx.doi.org/10.1007/109689873

20. Barski, A., Cuddapah, S., Cui, K., Roh, T.-Y., Schones, D.E., Wang, Z., Wei, G., Chepelev, I., Zhao, K.: High-resolution profiling of histone methylations in the human genome. Cell 129(4), 823–37 (2007). doi: 10.1016/j.cell.2007.05.009

21. Li, H., Durbin, R.: Fast and accurate long-read alignment with Burrows-Wheeler transform. Bioinformatics (Oxford, England) 26(5), 589–95 (2010). doi: 10.1093/bioinformatics/btp698

22. Langmead, B., Salzberg, S.L.: Fast gapped-read alignment with Bowtie 2. Nat Methods 9(4), 357–359 (2012). doi: 10.1038/nmeth.1923.#14603

23. Hoffmann, S., Otto, C., Kurtz, S., Sharma, C.M., Khaitovich, P., Vogel, J., Stadler, P.F., Hackermüller, J.: Fast mapping of short sequences with mismatches, insertions and deletions using index structures. PLoS computational biology 5(9), 1000502 (2009). doi: 10.1371/journal.pcbi.1000502

24. Kim, D., Pertea, G., Trapnell, C., Pimentel, H., Kelley, R., Salzberg, S.L.: TopHat2: accurate alignment of transcriptomes in the presence of insertions, deletions and gene fusions. Genome biology 14(4), 36 (2013). doi: 10.1186/gb-2013-14-4-r36

25. Hoffmann, S., Otto, C., Doose, G., Tanzer, A., Langenberger, D., Christ, S., Kunz, M., Holdt, L.M., Teupser, D., Hackermüller, J., Stadler, P.F.: A multi-split mapping algorithm for circular RNA, splicing, trans-splicing and fusion detection. Genome biology 15(2), 34 (2014). doi: 10.1186/gb-2014-15-2-r34

26. Trapnell, C., Williams, B.a., Pertea, G., Mortazavi, A., Kwan, G., van Baren, M.J., Salzberg, S.L., Wold, B.J., Pachter, L.: Transcript assembly and abundance estimation from RNA-Seq reveals thousands of new transcripts and switching among isoforms. Nature Biotechnology 28(5), 511–515 (2011). doi: 10.1038/nbt.1621.Transcript.171

27. Anders, S., Pyl, P.T., Huber, W.: HTSeq-A Python framework to work with high-throughput sequencing data. Bioinformatics 31(2), 166–169 (2015). doi: 10.1093/bioinformatics/btu638

28. Zhang, Y., Liu, T., Meyer, C.a., Eeckhoute, J., Johnson, D.S., Bernstein, B.E., Nusbaum, C., Myers, R.M., Brown, M., Li, W., Liu, X.S.: Model-based analysis of ChIP-Seq (MACS). Genome biology 9(9), 137 (2008). doi: 10.1186/gb-2008-9-9-r137

29. Fidler, F., Gordon, A.: Science is in a reproducibility crisis – how do we resolve it? The Conversation (2013). Available at: http://theconversation.com/science-is-in-a-reproducibility-crisis-how-do-we-resolve-it-16998

30. Lehrer, J.: The truth wears off. New Yorker Dec 13, 52–57 (2010)

31. Van Bavel, J.: Why do so many studies fail to replicate? The New York Times, 10 (2016). Available at: https://www.nytimes.com/2016/05/29/opinion/sunday/why-do-so-many-studies-fail-to-replicate.html

32. Fomel, S., Claerbout, J.F.: Guest Editors’ Introduction: Reproducible Research. Computing in Science & Engineering 11(1), 5–7 (2009). doi: 10.1109/MCSE.2009.14

33. Leisch, F.: Sweave: Dynamic generation of statistical reports using literate data analysis. In: Härdle, W., Rönz, B. (eds.) Compstat 2002 — Proceedings in Computational Statistics, pp. 575–580. Physica Verlag, Heidelberg (2002). ISBN 3-7908-1517-9. http://www.stat.uni-muenchen.de/texttildelowleisch/Sweave

34. Xie, Y.: knitr: A comprehensive tool for reproducible research in R. In: Stodden, V., Leisch, F., Peng, R.D. (eds.) Implementing Reproducible Computational Research. Chapman and Hall/CRC, Boca Raton, Florida (2014). ISBN 978-1466561595. http://www.crcpress.com/product/isbn/9781466561595

35. Pérez, F., Granger, B.E.: IPython: a system for interactive scientific computing. Computing in Science and Engineering 9(3), 21–29 (2007). doi: 10.1109/MCSE.2007.53

36. Afgan, E., Baker, D., Batut, B., van den Beek, M., Bouvier, D., Cech, M., Chilton, J., Clements, D., Coraor, N., Grüning, B.A., Guerler, A., Hillman-Jackson, J., Hiltemann, S., Jalili, V., Rasche, H., Soranzo, N., Goecks, J., Taylor, J., Nekrutenko, A., Blankenberg, D.: The galaxy platform for accessible, reproducible and collaborative biomedical analyses: 2018 update. Nucleic acids research 46, 537–544 (2018). doi: 10.1093/nar/gky379

37. Cingolani, P., Sladek, R., Blanchette, M.: BigDataScript: a scripting language for data pipelines. Bioinformatics 31(1), 10–16 (2014). doi: 10.1093/bioinformatics/btu595

38. Ewels, P., Krueger, F., Käller, M., Andrews, S.: Cluster flow: A user-friendly bioinformatics workflow tool. F1000Research 5, 2824 (2016). doi: 10.12688/f1000research.10335.2

39. Gafni, E., Luquette, L.J., Lancaster, A.K., Hawkins, J.B., Jung, J.-Y., Souilmi, Y., Wall, D.P., Tonellato, P.J.: Cosmos: Python library for massively parallel workflows. Bioinformatics 30, 2956–2958 (2014). doi: 10.1093/bioinformatics/btu385

40. Kaushik, G., Ivkovic, S., Simonovic, J., Tijanic, N., Davis-Dusenbery, B., Kural, D.: Rabix: An open-source workflow executor supporting recomputability and interoperability of workflow descriptions. Pacific Symposium on Biocomputing. Pacific Symposium on Biocomputing 22, 154–165 (2017). doi: 10.1142/9789813207813_0016

