## Supplemental Text and Figures for "uap: Reproducible and Robust HTS Data Analysis"

### Supplemental material: uap: Reproducible and Robust HTS Data Analysis

June 24, 2019

#### List of Figures

Figure S1 DAG of the analysis of ChIP-seq data from Barski *et al.*. . . . . . . . 4
Figure S2 Rendered graph of a finished `sam_to_sorted_bam` run. . . . . . . 5

#### List of Tables

Table S1 Comparison of workflow management systems based on essential features, particularly regarding reproducibility. . . . . . . . . . . . . . . . . . . . 2
Table S2 Availability of software tools used as pipeline and/or work-flow managers. 3

Table S1: **Comparison of workflow management systems based on essential features, particularly regarding reproducibility.** Tools were selected from `https://github.com/pditommaso/awesome-pipeline` retaining only tools that were considered to be actively developed by at least five contributors (latest commit after 31.01.2018), are licensed as open source, where the scope of application is clearly on bioinformatics, and which support cluster batch systems.
**Modular**: Workflows are assembled from reusable steps; **Flexible**: Configuration of analysis is separated from its code (code $\neq$ definition) and workflows can easily be adapted without programming skills; **Failure recovery**: Failure during execution is detected and analyses halted to avoid corrupted results; **Reproducibility - Dependencies**: WMS enforces that dependencies between analysis steps and intermediate results are correctly maintained. **Consistency**: WMS safeguards that analysis steps are successfully completed prior to execution of subsequent steps. **Linking code and result**: WMS ensures consistency between the code defining the analysis and the currently available results. **Logging**: WMS logs the mentioned information; **Data authenticity** WMS records information about creator and creating process(es) of the data; **Data integrity**: WMS enables validating the pristine state of created data, e.g. using hash sums; **Supp. platforms**: WMS is designed to work with these technologies; **Docker**: WMS can run jobs using Docker images **Supp. CWL**: WMS supports CWL;
**Locally executable**: WMS can be without cluster or (web)server; Results are given as ○ (not met), ◐ (partially met), ● (fully met) and – (not stated). In case of supported platforms, ● corresponds to running out of the box, while ◐ indicates that the WMS can be used on the respective platform upon adaptataions of the configuration or extension of code. Ratings are based on information provided in the papers, documentations and manuals. Details on the availability of these tools are provided in Supplemental Table S2.

| | Arvados | bcbio-nextgen[1] | BigDataScript[2] | Bpipe[3] | Cluster Flow | Cosmos | Cromwell | Galaxy | Gwf | Hive | Nextflow | PypeFlow | Rabix | Toil | uap |
|---|---|---|---|---|---|---|---|---|---|---|---|---|---|---|---|
| Modular | ◐ | ○ | ◐ | ● | ● | ◐ | ● | ● | ◐ | ● | ◐ | – | ◐ | ◐ | ● |
| Flexible | ● | ● | ○ | ◐ | ● | ○ | ● | ● | ○ | ◐ | ● | – | ● | ● | ● |
| Failure recovery | – | – | ● | ● | – | ● | ● | ● | – | – | ● | – | ◐ | ● | ● |
| Reproducibility | | | | | | | | | | | | | | | |
| Dependencies | – | – | ◐ | ◐ | – | ○ | ● | ● | – | – | ● | – | – | – | ● |
| Consistency | ● | ● | ● | ● | ● | ● | ● | ● | ● | ● | ● | ● | ● | ● | ● |
| Linking code and result | – | – | – | – | – | – | – | ● | – | – | ● | – | – | – | ● |
| Logging | | | | | | | | | | | | | | | |
| Stdout/Stderr | – | ● | ● | ● | ● | – | ● | ● | ◐ | – | ◐ | – | – | ◐ | ● |
| Exit status | ● | – | ● | ● | – | – | ● | ● | – | – | ● | – | – | – | ● |
| Tool versions | – | – | – | ◐ | ◐ | – | – | ● | – | – | – | – | – | – | ● |
| WMS version | ● | – | – | – | – | – | – | – | – | – | ● | – | – | – | ● |
| Executed commands | ● | ● | ● | ● | ◐ | ● | – | ● | – | – | ● | – | – | – | ● |
| Execution date | ● | – | – | ● | – | ● | – | ● | – | ○ | ● | – | – | – | ● |
| In-/output files | ● | – | – | ● | ● | ◐ | – | ● | – | – | – | – | – | – | ● |
| Data authenticity | ● | – | – | – | – | ● | – | ● | – | – | – | – | – | – | ● |
| Data integrity | ● | – | – | – | ◐ | ○ | – | ○ | – | – | – | – | ● | – | ● |
| Supp. platforms | | | | | | | | | | | | | | | |
| SGE/OGE/UGE | ○ | ● | ● | ● | ● | – | ● | ● | ● | ● | ● | ◐ | ◐ | ● | ● |
| SLURM | ● | ● | ● | ○ | ● | ● | ● | ● | ● | ○ | ● | ◐ | ◐ | ● | ● |
| Torque | ○ | ● | ○ | ○ | ○ | – | ◐ | ● | ○ | ● | ● | ◐ | ◐ | ● | ◐ |
| Moab | ○ | – | ● | ○ | ○ | – | ◐ | ○ | ○ | ○ | ○ | ○ | ◐ | – | ◐ |
| LSF | ○ | ● | ◐ | ● | ● | ● | ● | ● | ○ | ● | ● | ◐ | ◐ | ● | ◐ |
| Cloud (EC2) | ◐ | ◐ | ◐ | – | – | ● | ◐ | ● | – | ◐ | ● | – | ● | ● | ○ |
| Docker | ● | ● | – | – | – | – | ● | ● | – | ● | ● | – | ● | ● | ○ |
| Supp. CWL | ● | ◐ | – | ○ | ○ | ○ | ● | ● | ○ | ○ | ● | ○ | ● | ● | ○ |
| Locally executable | ○ | ● | ● | ● | ● | ● | ● | ◐ | ● | ● | ● | – | ● | ● | ● |

Table S2: **Availability of software tools used as pipeline and/or work-flow managers.**

| Tool | Reference | url |
|---|---|---|
| Arvados | | https://arvados.org/ |
| bcbio-nextgen | [1] | https://github.com/bcbio/bcbio-nextgen |
| BigDataScript | [2] | https://pcingola.github.io/BigDataScript/ |
| Bpipe | [3] | https://github.com/ssadedin/bpipe |
| Cluster Flow | [4] | http://clusterflow.io |
| Cosmos | [5] | https://cosmos.hms.harvard.edu |
| Cromwell | | https://github.com/broadinstitute/cromwell |
| Galaxy | [6] | https://usegalaxy.org/ |
| Gwf | | https://github.com/gwforg/gwf |
| Hive | | https://github.com/Ensembl/ensembl-hive |
| Nextflow | [7] | https://www.nextflow.io/ |
| PypeFlow | | https://github.com/PacificBiosciences/pypeFLOW |
| Rabix | [8] | https://github.com/rabix/rabix |
| Toil | | https://github.com/DataBiosphere/toil |
| uap | | https://github.com/yigbt/uap |

Figure S1: **DAG of the analysis of ChIP-seq data from Barski *et al.* [9]**. Yellow boxes contain the names of the steps and, if step name and type differ, additionally the type of the step in brackets. The adjacent, attached boxes display the number of finished versus total runs. The color coding of the latter boxes indicates whether that step has been finished (green), partially finished (ocher), or not started yet (red).

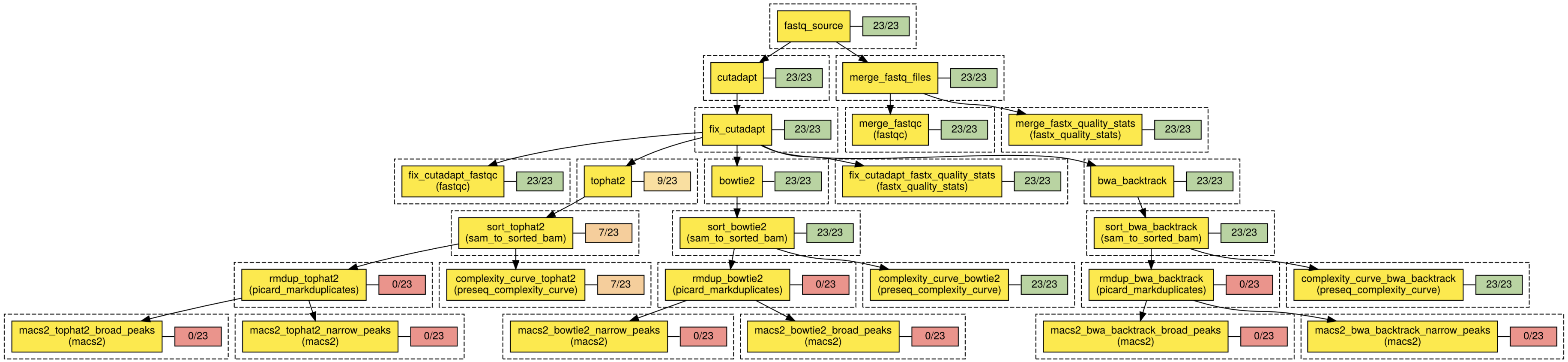

Figure S2: **Rendered graph of a finished `sam_to_sorted_bam` run.** The graph was created by `uap`'s `render` subcommand and manually edited to better fit to the page. It shows CPU and memory usage of the run and its executed commands.

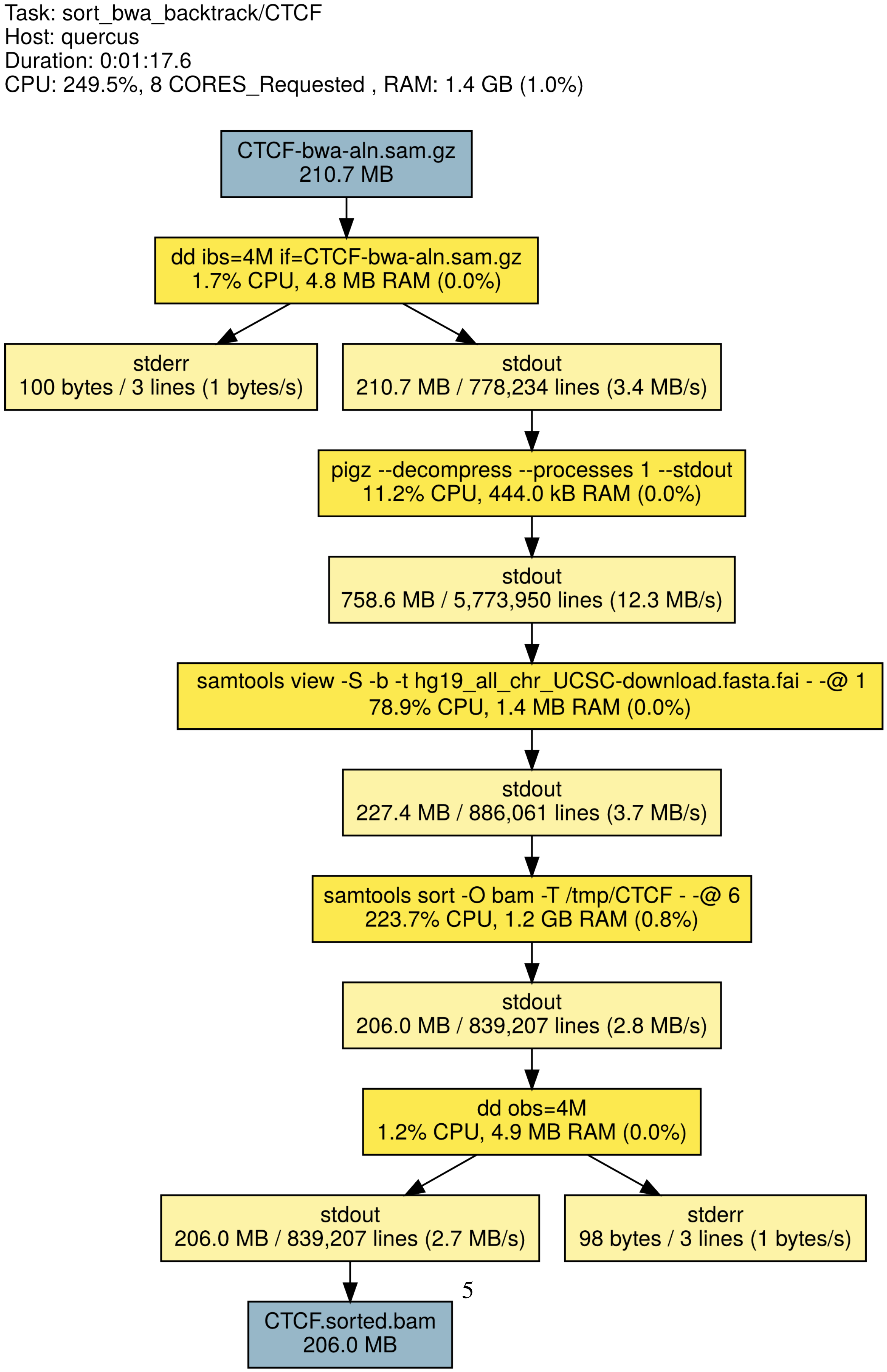

### References


[1] Guimera, R.V.: bcbio-nextgen: Automated, distributed next-gen sequencing pipeline. EMBnet.journal **17**, 30 (2012). doi:10.14806/ej.17.B.286

[2] Cingolani, P., Sladek, R., Blanchette, M.: BigDataScript: a scripting language for data pipelines. Bioinformatics **31**(1), 10–16 (2014). doi:10.1093/bioinformatics/btu595

[3] Sadedin, S.P., Pope, B., Oshlack, A.: Bpipe: a tool for running and managing bioinformatics pipelines. Bioinformatics (Oxford, England) **28**(11), 1525–6 (2012). doi:10.1093/bioinformatics/bts167

[4] Ewels, P., Krueger, F., Kller, M., Andrews, S.: Cluster flow: A user-friendly bioinformatics workflow tool. F1000Research **5**, 2824 (2016). doi:10.12688/f1000research.10335.2

[5] Gafni, E., Luquette, L.J., Lancaster, A.K., Hawkins, J.B., Jung, J.-Y., Souilmi, Y., Wall, D.P., Tonellato, P.J.: Cosmos: Python library for massively parallel workflows. Bioinformatics **30**, 2956–2958 (2014). doi:10.1093/bioinformatics/btu385

[6] Afgan, E., Baker, D., Batut, B., van den Beek, M., Bouvier, D., Cech, M., Chilton, J., Clements, D., Coraor, N., Grning, B.A., Guerler, A., Hillman-Jackson, J., Hiltemann, S., Jalili, V., Rasche, H., Soranzo, N., Goecks, J., Taylor, J., Nekrutenko, A., Blankenberg, D.: The galaxy platform for accessible, reproducible and collaborative biomedical analyses: 2018 update. Nucleic acids research **46**, 537–544 (2018). doi:10.1093/nar/gky379

[7] Di Tommaso, P., Chatzou, M., Floden, E.W., Barja, P.P., Palumbo, E., Notredame, C.: Nextflow enables reproducible computational workflows. Nature biotechnology **35**, 316–319 (2017). doi:10.1038/nbt.3820

[8] Kaushik, G., Ivkovic, S., Simonovic, J., Tijanic, N., Davis-Dusenbery, B., Kural, D.: Rabix: An open-source workflow executor supporting recomputability and interoperability of workflow descriptions. Pacific Symposium on Biocomputing. Pacific Symposium on Biocomputing **22**, 154–165 (2017). doi:10.1142/9789813207813_0016

[9] Barski, A., Cuddapah, S., Cui, K., Roh, T.-Y., Schones, D.E., Wang, Z., Wei, G., Chepelev, I., Zhao, K.: High-resolution profiling of histone methylations in the human genome. Cell **129**(4), 823–37 (2007). doi:10.1016/j.cell.2007.05.009
